# Experimental Urolithiasis Model to assess Phyto-fractions as Anti-lithiatic Contributors: A Herbaceutical Approach

**DOI:** 10.1101/2021.06.01.446538

**Authors:** Aishwarya Tripurasundari Devi, N Yashaswini, Farhan Zameer, MN Nagendra Prasad

**Affiliations:** Department of Biotechnology, JSS Science and Technology University, JSS Research Foundation, SJCE Campus, Manasagangotri, Mysore - 560 006, Karnataka, India; PathoGutOmic Laboratory, Department of Biochemistry, School of Basic and Applied Sciences, Dayananda Sagar University, Shavige Malleshwara Hills, Kumaraswamy Layout, Bengaluru - 560 111, Karnataka, India

**Author notes:** Both the authors share the corresponding authorship for equal contribution (Farhan Zameer, MN Nagendra Prasad). **Corresponding author 1: Dr. Nagendra Prasad MN**: Professor & Head, Department of Biotechnology, JSS Science and Technology University, JSS Research Foundation, SJCE Campus, Manasagangotri, Mysore - 560 006, Karnataka, India., **Corresponding author 2: Dr. Farhan Zameer:** Assistant Professor in Biochemistry, PathoGutOmic Laboratory, School of Basic and Applied Sciences, Department of Biological Sciences, Dayananda Sagar University, Shavige Malleshwara Hills, Kumaraswamy Layout, Bengaluru - 560 111, Karnataka, India.

**Keywords:** Lithiasis, Kidney stones, Calcium oxalate, Biomarkers, Anti-stress, Structure-activity relationship (SAR) Studies, Botanicals, *Kalanchoe pinnata*, Phyto-pharmaceutics

## Abstract

Life-style disorders have bought a serious burden on the maintenance of health in animals and humans. Lithiasis specifically nephro- and urolithiasis is no exception and needs urgent attention. Currently, only semi-invasive and surgical methods are widely employed which leads to trauma and reoccurrence of kidney stones. Hence complementary and alternative herbal medicine could pave newer ways in exploring anti-lithiatic contributors. The current study attempts to screen twenty herbal hot aqueous leaf extracts for assessing their antioxidant potency (anti-stress) and efficiency against urolithiasis in an experimental calcium oxalate-induced *in vitro* (chicken egg membrane) model. The study was further validated by *In silico* molecular docking studies using the Molegro software package on enzymatic biomarkers involved in scavenging oxidants in the host and regulating oxalate metabolism at a cellular level. Among the screened botanicals *Kalanchoe pinnata* exhibited promising results compared to the standard chemical (potassium-magnesium citrate) and phyto-formulation drug (cystone) currently used by clinicians for treating urolithiasis. The phytochemical profiling (qualitative and quantitative) and virtual studies indicated rutin from *Kalanchoe pinnata* as a potential candidate for preventing kidney stones. The results of the current study provide better insights into the design and development of newer, smart, and cost-effective herbal therapeutics making food as medicine.

**Graphical Abstract:** 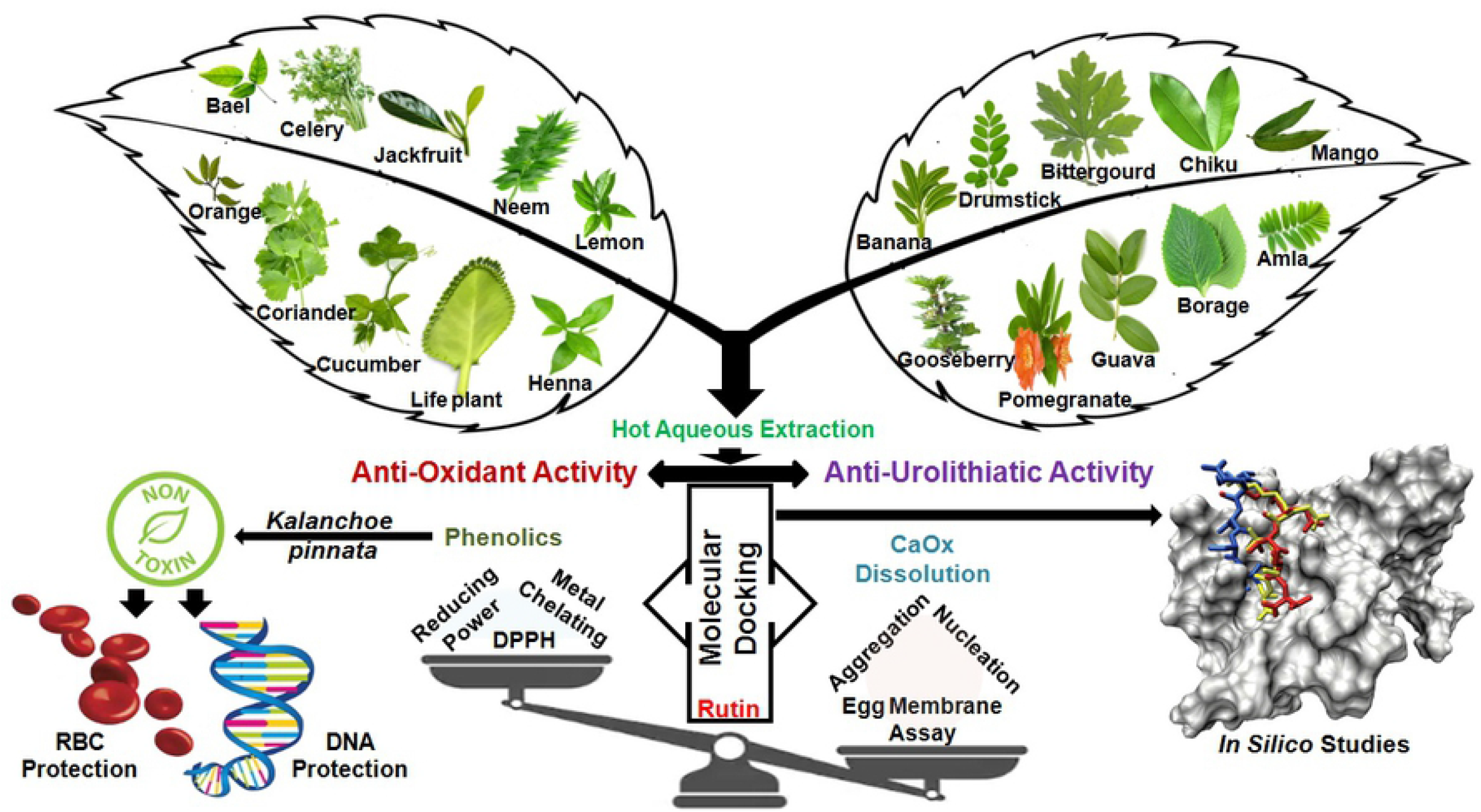

## Introduction

Food and health are the two integral components of human wellbeing. However, in the urge of adapting to newer technologies and advanced niches both these components are highly neglected. In consequence, many lifestyle, non-communicative disorders, or diseases has become a part of human health. Indian folklore medicine and Ayurveda proposes the implications of “food as medicine” from time immemorial. Further, complementary and alternative medicine (CAM) has been successful in exploring the scientific proof and mechanism of actions of these phyto-formulations leading to the development of newer drug targets and therapeutics. Phyto-cocktails are effective in many complications from venom antidots (Janardhan et al., 2019; Vineetha et al., 2020, Bhavya et al., 2021), neurodegenerative disorders (Kunnel et al., 2019; Satapathy et al., 2020), diabetes (Putta et al., 2016), ulcers (Prasad et al., 2019), cancer (Rakesh et al., 2015), infection (Zameer et al., 2016) to inflammation (Rakesh et al., 2016). With all the set examples from our previous studies, one major disorder caught our attention which was a common lifestyle disorder yet very painful, which had no structured therapy and was found in folks of all ages but most prevalent in men known as lithiasis (stone formation).

Lithiasis is a condition in which the formation of stones or calculi is observed due to the concentration of mineral salts (Grases et al., 2007). The kidney, urinary tract, pancreas, and gallbladder are the most commonly affected organs by lithiasis, and they are categorized based on their site of formation leading to nephrolithiasis, urolithiasis, pancreatolithiasis, and cholecystolithiasis respectively (Tatapudi et al., 2020). Stones containing calcium are a common type of calculi with a prevalence of 70-80%, of which calcium phosphate and calcium oxalate dominate (Han et al., 2015). Supersaturation of urine in presence of calcium and oxalate facilitates calcium oxalate stone development (Paliouras et al., 2012). Following are the factors which affect the formation of calcium oxalate stones; acidic urine, low volume of urine, hypercalciuria (increased concentration of calcium in urine leads to precipitation of calcium salts), hyperoxaluria (high concentration of oxalate excreted in urine), hypocitraturia (excretion of a lower amount of citrate in urine which leads to high pH) and hyperuricosuria (acidic urine dissolves uric acid, leading to stone formation). Apart from calcium, magnesium phosphate, uric acid, cysteine, silica, xanthine, and 2, 8-dihydroxyadenine also account for the stone formation (Moe et al., 2006; Evan et al., 2015). Further, with all the above understanding of biochemistry and pathophysiology of urolithiasis, an optimized and standardized model system is a prime necessity that should be rapid, reliable, and reproducible. To serve this purpose, the chicken egg membrane model was employed to study the dissolution of calcium oxalate with nucleation and aggregation assessments (Phatak and Hendre, 2015). Indian traditional system of medicine (Ayurveda) strongly emphasizes the use of phyto-formulations for almost any ailments and urolithiasis, is no exception. Hence an attempt was made to explore the less exploited plants as anti-urolithiatic agents (table 1).

**Table 1:** List of selected 20 botanicals screened for anti-oxidant and anti-urolithiatic activity.

| SL No | BOTANICAL NAME | COMMON NAME | FAMILY | BIOLOGICAL ACTIVITY | REFERENCES |
| --- | --- | --- | --- | --- | --- |
| 1 | *Aegle marmelos* | Bael | *Rutaceae* | Antioxidant and anti-inflammatory properties | Pynam & Dharmesh (2018) |
| 2 | *Apium graveolens* | Celery | *Apiaceae* | Antioxidant, neurochemical activity and also used in treatments for hypertension, gout, and diabetes | Chonpathompikunlert *et al.*, (2018) |
| 3 | *Artocarpus heterophyllus* | Jackfruit | *Moraceae* | Anticarcinogenic, antimicrobial, antifungal, anti-inflammatory, wound healing, and hypoglycemic effects | Ranasinghe *et al.*, (2019) |
| 4 | *Azadirachta indica* | Neem | *Meliaceae* | Hypolipidemic, antifertility, microbicidal, antidiabetic, anti-inflammatory, hepatoprotective, antipyretic, hypoglycemic, insecticidal, nematicidal, antiulcer, antioxidant, neuroprotective, cardioprotective, and antileishmaniasis properties. | Saleem *et al.*, (2018) |
| 5 | *Citrus limon* | Lemon | *Rutaceae* | Antiuro lithiatic activity | Prabhu *et al.*, (2016) |
| 6 | *Citrus sinensis* | Orange | *Rutaceae* | Antioxidant, Antiuro lithiatic activity | Ungjaroenwathana *et al.*, (2019) |
| 7 | *Coriandrum sativum* | Coriander | *Apiaceae* | Antimicrobial, antioxidant, hypoglycemic, hypolipidemic, anxiolytic, analgesic, anti-inflammatory, anti-convulsant and anticancer activities | Laribi *et al.*, (2015) |
| 8 | *Cucumis sativus* | Cucumber | *Cucurbitaceae* | Anti-inflammatory, antioxidant activity | Trejo-Moreno *et al.*, (2018) |
| 9 | *Kalanchoe pinnata* | Life plant | *Crassulaceae* | Antilithiatic activity, antinociceptive, antiedematogenic, and anti-inflammatory | Sohgaura *et al.*, (2018)<br>Ferreira *et al.*, (2014) |
| 10 | *Lawsonia inermis* | Henna | *Lythraceae* | Antibacterial, antifungal, antioxidant, antiproliferative, cytotoxicity | Elansary *et al.*, (2020) |
| 11 | *Mangifera indica* | Mango | *Anacardiaceae* | Antioxidant, anti-inflammatory, radioprotective, antitumor, immune-modulatory, antiallergic, antidiabetic, anti-bone resorption, mono-amine oxidase inhibiting, antiviral, antifungal, antibacterial, antispasmodic, antidiarrheal, antimalarial, antiparasitic lipolytic properties. | Batool *et al.*, (2018) |
| 12 | *Manilkara zapota* | Chiku | *Sapotaceae* | Antibacterial, hepato-protective, anti-inflammatory, anti-tussive, antifungal, anti-tumour, and free radical scavenging potential | Sharma *et al.*, (2020) |
| 13 | *Momordica charantia* | Bitter gourd | *Cucurbitaceae* | Antihyperglycemic, antibacterial, antiviral, antitumor, immunomodulation, antioxidant, antidiabetic, anthelmintic, antimutagenic, antiulcer, antilipolytic, antifertility, hepatoprotective, anticancer and anti-inflammatory activities | Jia *et al.*, (2017) |
| 14 | *Moringa oleifera* | Drumsticks | *Moringaceae* | Antiuro lithiatic activity, analgesic, anti-inflammatory, antipyretic, anticancer, antioxidant, nootropic, hepatoprotective, astroprotective, antiulcer, cardiovascular, antiobesity, antiepileptic, antiasthmatic, antidiabetic, antiuro lithiatic, diuretic, local anesthetic, antiallergic, anthelmintic, wound healing, antimicrobial, immunomodulatory, and antidiarrheal properties | Karadi *et al.*, (2006)<br>Bhattacharya *et al.*, (2018) |
| 15 | *Musa paradisiaca* | banana | *Musaceae* | Antiuro lithiatic, antioxidant activity | Panigrahi PN *et al.*, (2017) |
| 16 | *Phyllanthus emblica* | Indian gooseberry or amla | *Euphorbiaceae* | Antidiabetic, hypolipidemic, antibacterial, antioxidant, antiulcerogenic, hepatoprotective, gastroprotective, and chemopreventive properties. In treatment of diarrhea, jaundice, and inflammation. | Krishnaveni & Mirunalini (2010)<br>Li *et al.*, (2020) |
| 17 | *Plectranthus amboinicus* | Country Borage | *Lamiaceae* | Antimicrobial, anti inflammatory, antitumor, wound healing, anti-epileptic, larvicidal, antioxidant and analgesic activities. Effective against respiratory, cardiovascular, oral, skin, digestive and urinary diseases | Arumugam *et al.*, (2016) |
| 18 | *Psidium guajava* | Guava | *Myrtaceae* | Antiuro lithiatic activity, antioxidant, anti inflammatory activity | Agarwal & Varma (2015)<br>Vasconcelos *et al.*, (2017) |
| 19 | *Punica granatum* | Pomegranate | *Lythraceae* | Antiuro lithiatic activity, anti-inflammatory, antibacterial, antidiarrheal, immune modulatory, antitumor, wound healing and antifungal | Rathod *et al.*, (2012)<br>Saeed *et al.*, (2018) |
| 20 | *Ribes Uva-crispa* | Gooseberry | *Grossulariaceae* | Antioxidant activity | Laczko-Zöld *et al.*, (2018) |

Further, phytobioactives in different forms crude, partially purified, purified, or in cocktails play a vital role either as a single drug or complex provide synergy in ameliorating health complications from a simple infection, inflammation to a cascade of disorders/diseases (Aishwarya et al., 2020). These various activities are mainly due to the different classes of phytochemicals predominately secondary metabolites of polyphenolic umbrella namely phenolics, flavonoids, alkaloids, terpenoids, tannins, saponins, and sterols (Pankaj et al., 2020; Khan et al., 2020). According to Gulcin (2020), a detailed correlation of phytochemicals with antioxidant efficacy is well illustrated and this property could be explored for stress and inflammation which is usually encountered during urolithiasis. Apart from the antioxidant phytobioactives, enzymatic biomarkers such as catalase (CAT), superoxide dismutase (SOD), peroxidase (PER), glutathione S-transferase (GST), contribute against redox reaction, stress, and inflammation respectively (Gopal et al., 2009). Besides, metabolic enzymes such as alanine-glyoxlate aminotransferase, oxalyl-coA decarboxylase, D-glycerate dehydrogenase, and lactate dehydrogenase (LDH) regulate a crucial role in stone formation (Ramu et al., 2017). Elucidation of the mechanism of action of these biomarkers in urolithiasis is very much essential. Structure-activity relationship (SAR) studies were employed to decipher the above set objective using molecular docking studies (Madhusudhan et al., 2016).

Henceforth, in the current study, an attempt has been employed to profile phytochemicals (qualitative), to evaluate the antioxidant potency, and to adapt a simple experimental model for kidney stones (chicken egg membrane) to screen anti-urolithiatic effect (*in vitro*) of selected 20 plants (aqueous extracts) supported by the *in silico* evaluation for proof-of-concept in search of highly-efficient and cost-effective herbaceuticals to ameliorate urolithiasis.

## Materials & Methods

All chemicals, solvents, and reagents used for the study were of analytical grade and were purchased from Himedia Pvt. Ltd, Mumbai, India. The standard drug Cystone^®^ was purchased from Himalaya Drug Company, Bangalore, India. Unfertilized chicken eggs were procured from local poultry in Mysuru and membranes were isolated.

### Plant material

Twenty different medicinal plant leaves (table 1) were collected from in and around Mysuru, India, from March to July 2018. Further, the plants were authenticated and are deposited at the herbarium center, Department of Studies in Botany, University of Mysore, Karnataka. The leaves of *Aegle marmelos, Apium graveolens, Artocarpus heterophyllus, Azadirachta indica, Citrus limon, Citrus sinensis, Coriandrum sativum, Cucumis sativus, Kalanchoe pinnata, Lawsonia inermis, Mangifera indica, Manilkara zapota, Momordica charantia, Moringa oleifera, Musa paradisiaca, Phyllanthus emblica, Plectranthus amboinicus, Psidium guajava, Punica granatum* and *Ribes Uva-crispa* were harvested, leaves were surface sterilized and dried in a hot air oven at 45°C for 48 hours and powdered to 60 mesh packed in an air-tight container until further experimentation.

### For extraction

The hot aqueous extract was prepared for all the twenty plants, 20g of finely powdered leaves of the individual plant were mixed with 100 mL of distilled water (pH 5.8) and kept over a magnetic stirrer for 3 hours with a constant heat around 50 ± 2°C (Meghashri et al., 2011). Only aqueous extracts were targeted in the study, intending to isolate polar/hydrophilic phyto-molecules which will be much more practical to suggest patients with urological complications as a homemade decoction. Muslin cloth was used to filter; later, the filtrate was subjected to centrifugation for 10 min at 8000 rpm, after which the supernatant was collected and stored in a freezer for further use.

### Phytochemical analysis

All the qualitative phytochemical profiling was performed according to the AOAC method (Shameh et al., 2018) and is represented in supplementary table 1.

### Estimation of total phenol content (TPC)

All aqueous plant extracts were assessed by Folin-Ciocalteau (FC) method with a slight modification (Meghashri et al., 2010). Briefly, various volumes of the extracts were made up to 3 mL with distilled water. Folin-Ciocalteau reagent of (1:1 diluted) 0.5 mL was added and incubated for 5 min at room temperature, 1.5 mL of sodium carbonate (20%) was added followed by incubation in a boiling water bath for 5 min. The absorbance was measured at 640 nm. Gallic acid (1mg/mL) equivalents were used as the standard to estimate the phenolic content.

### Assessment of *In vitro* antioxidant activities for aqueous phyto-fractions

#### DPPH free radical scavenging assay

The free radical scavenging activity of all the plant extracts was measured by the DPPH (1,1-diphenyl-2picrylhydrazyl) method (Meghashri et al., 2010). Varied concentrations of the extracts [5 μg/mL, 10 μg/mL, 15 μg/mL, 20 μg/mL, and 25 μg/mL of gallic acid equivalent] followed by 1 mL of DPPH solution, shaken and was incubated at room temperature for 20 min in the dark after which absorbance was measured at 517nm. Gallic acid was used as standard. The activity of the radical scavenging was calculated by the following equation:

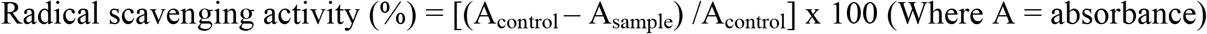

#### Metal ion chelating assay

The complex ferrous iron - ferrozine can chelate Fe^2+,^ which can be observed at 562 nm (Meghashri et al., 2010). Different concentrations of all extracts were taken, 50 μL of FeCl_2_ and 200 μL of ferrozine were mixed and incubated at room temperature in the dark for 10 min; finally, absorbance at 562 nm against blank was measured. The same equation used in DPPH scavenging activity is used here to know the capacity of the extract in cheating ferrous ions with EDTA considered as standard.

#### Reducing power assay

Reduction of iron (III) by the extracts was measured with slight modification (Nedamani et al., 2015) by adding the various concentration of all extracts and made up the volume to 500 μL with phosphate buffer (20 mM) and then 500 μL of sodium ferricyanide (1%) was added and incubated at 50°C for 20 min. 500 μL of trichloroacetic acid (10%) was added to the solution to terminate the reaction followed by 10 min centrifugation. Finally, 1.5 mL of distilled water along with 300 μL of ferric chloride (0.1%) was added to the supernatant, and absorbance at 700nm was measured using gallic acid as standard.

#### Anti-lithiatic activity

The anti-lithiatic activity was assessed for all the extracts according to Phatak and Hendre, (2015). Calcium chloride (25 mM) 1mL of the solution was mixed with 2 mL of tris buffer (pH 7.4). Water was considered as control, and extracts were added to the above mixture; finally, in the end, 1 mL of sodium oxalate (25 mM) was added, and the clock was started. Absorbance at 620 nm for 10 min was measured and a standard graph.

#### Preparation of semi-permeable membrane from chicken eggs

Among the 20 extracts screened, only the top five extracts (*Kalanchoe pinnata*, *Musa paradisiaca, Punica granatum, Coriandrum sativum,* and *Moringa oleifera)* were used for further *in vitro* anti-lithiatic egg membrane assay. The complete content of the eggs was removed by piercing with a glass rod on top of the eggs. Then these eggs were washed carefully with distilled water and were immersed in a beaker containing 2N HCL overnight for decalcification. The following day the semi-permeable membrane was cautiously removed from the shells rinsed with distilled water and neutralized with ammonia solution to remove any traces of acid from the membranes and finally rinsed with distilled water. Further, it was refrigerated at pH 7-7.4 condition to maintain moisture (Phatak and Hendre, 2015).

#### Synthesis of calcium oxalate through homogenous precipitation

Calcium chloride dihydrate weighing 1.47g was dissolved using 100mL of distilled water, and 100mL of 2N H_2_SO_4_ was used to dissolve 1.34g of sodium oxalate. Calcium oxalate starts precipitating with constant stirring when both the above solutions are mixed. Traces of sulphuric acid is removed by rinsing with ammonia solution and then finally rinsed with distilled water and kept for drying at 60°C for 4 hours (Monika et al., 2012). Calcium oxalate 1 mg/mL served as a negative control and a Cystone concentration of 20 mg/mL was set as a positive control (Phatak and Hendre, 2015). Further, the experimental groups were as follows, Negative control: 1 mL calcium oxalate + 1mL of distilled water, Positive control: 1 mL of calcium oxalate + 1 mL of Cystone solution, *Kalanchoe pinnata*: 1 mL of calcium oxalate + 500 μL of extract, *Punica granatum:* 1 mL of calcium oxalate + 500 μL of extract, *Musa paradisiaca:* 1 mL of calcium oxalate + 500 μL of extract, *Coriandrum sativum:* 1 mL of calcium oxalate + 500 μL of extract and *Moringa oleifera:* 1 mL of calcium oxalate + 500 μL of extract respectively.

The above-mentioned groups consisting of both positive and negative controls along with extracts were packed into respective semi-permeable membranes further the mouth of the membrane was tied with the help of thread and was immersed in a conical flask containing 100 mL of 0.1 M tris buffer. Incubator was preheated for 2 hours to obtain a constant temperature of 37°C; all the groups were placed inside the incubator for 7-8 hours (Phatak and Hendre, 2015). The contents of each group were transferred into a fresh flask, and 2 mL of 1N sulphuric acid was added and was titrated against 0.9494 N KMnO_4_ till the endpoint that is light pink color is observed. Percentage dissolution was calculated based on titration (Monika et al., 2012). Based on these results the best aqueous extract exhibiting anti-lithiatic activity was found to be *Kalanchoe pinnata* and further studies focused on this single phyto-fraction only.

#### Standard curve of calcium oxalate

Potassium permanganate 3.2 g was dissolved in 1000 mL of distilled water and then boiled for 30 min. Later, filtered using Whatman filter paper to obtain 0.02M of KMnO_4_. Different concentrations of calcium oxalate (0.2 mg/mL, 0.4 mg/mL, 0.6 mg/mL, 0.8 mg/mL, and 1 mg/mL) was used and the volume to 1 mL using distilled water. A 4 mL of sulphuric acid was added with 80 μL of 0.02M KMnO_4_. The above mixture was mixed well and incubated for 2 hours after which absorbance was read using a spectrophotometer at 620 nm. For all further analyses, only the best extract (*Kalanchoe pinnata*) was intervened to deduce the probable mechanism of calcium oxalate dissolution.

#### Nucleation assay

The nucleation of calcium oxalate crystals was estimated using a spectrophotometer, and *Kalanchoe pinnata* inhibiting potency was determined by the method of Saha and Verma (2013), with minor alteration. Calcium chloride 4 mmol/L and sodium oxalate 50 mmol/L were mixed to initiate crystallization, and this was added to artificial urine. The solutions were prepared in Tris 0.05mol/L for calcium chloride and NaCl 0.15mol/L at pH 6.5 and 37°C for sodium oxalate. Nucleation rate was obtained by equating the time of crystal formation about the presence of varying concentrations of *Kalanchoe pinnata* and with no extract in another and also Cystone was used as the positive control, absorbance was recorded at 1 hour, 3 hours, and 24 hours at 620 nm, percentage inhibition was calculated accordingly.

#### Aggregation assay

Saha and Verma (2013) method was tailored slightly to obtain the rate of aggregation of crystals of calcium oxalate. Calcium chloride and sodium oxalate 50 mmol/L solutions were mixed to obtain COM crystals. The two solutions were equalized by incubating in a water bath at 60°C for 1 hour and then brought to 37°C followed by evaporation. These crystals were brought to a concentration of 1 mg/mL by dissolving in 0.05 mol/L Tris and 0.15 mol/L NaCl at 6.5 pH. Different concentrations of *Kalanchoe pinnata* were used along with negative control and positive control being Cystone were observed under a light microscope.

#### RBC protection assay by *Kalanchoe pinnata* extract

Consent from the healthy individual to procure erythrocytes was taken. The blood which was heparinized was centrifuged for 15 min at 1000g through which the buffy coat and plasma were separated, and the erythrocytes with the help of PBS were rinsed thrice at room temperature reintroduced into PBS for further analysis, and the volume was made up four times. For 5 min, *Kalanchoe pinnata* was incubated with erythrocytes and then further incubated for 1 hour at 37°C with hydrogen peroxide, ferric chloride, and ascorbic acid and kept in a shaker incubator for incubation and was observed under an optical microscope for any changes in the morphology (Beulah et al., 2015).

#### DNA protection assay by *Kalanchoe pinnata* extract

Protocol from Meghashri et al., (2010) with slight variations was adapted to check the efficacy of *Kalanchoe pinnata* in protecting the DNA was checked using lambda phage DNA (Meghashri et al., 2010). In the presence or lack of *Kalanchoe pinnata* and gallic acid, the oxidation with the help of Fenton’s reagent along with lambda DNA was estimated for 2 hours at 37°C. Electrophoresis was performed for the samples using 1% agarose gel at 50V DC for 2 hours and stained the gels using ethidium bromide and visualized using gel documentation.

#### High-Performance Liquid Chromatography (HPLC) Profiling

The crude aqueous fraction, a hot extract of *Kalanchoe pinnata* was subjected to HPLC in a C-18 column with different mobile phase water: acetonitrile in the ratio 80:20 (Meghashri et al., 2010). The following standards were used to measure Ferulic acid (321nm), Gallic acid (272nm), Rutin (257nm), Quercetin (257nm), and Vanillic acid (261nm) with 1mL/min flow rate.

#### Molecular Docking

*In silico* analyses provides molecular insights (Ranganatha et al., 2014; Satapathy et al., 2020) on the interaction of ligands (phyto-molecules) with that of the receptors (target enzymes). In the current study, two major classes of targets were analyzed (1). The enzymes regulating redox machinery and (2). Enzymes involved in calcium oxalate regulation. Henceforth, four major antioxidant enzymes namely Superoxide dismutase (PDB: 2V0A), Peroxidase (PDB: 1PRX), Catalase (PDB: 5GKN), and Glutathione S-Transferase (PDB: 1LJR) were retrieved from Protein Data Bank (PDB). This was backed up by metabolic enzymes Alanine-Glyoxylate aminotransferase (PDB: 1HOC), Oxalyl-Coa decarboxylase (PDB: 2JIB), D-glycerate dehydrogenase (PDB: 1GDH), and Lactate dehydrogenase (PDB: 5ZJF) were considered respectively. Four major ligand molecules were docked on the catalytic site of the receptors based on the HPLC profiling *i.e.,* Gallic acid (PubChem: 370), Vanillic acid (PubChem: 8468), Rutin (PubChem: 5280805), and Potassium Magnesium Citrate (PubChem: 10437763) among which the latter served as the standard drug which is currently used for treating kidney stones. Molegro Docking software package was used with visual docker and the results were analyzed considering the atomic contact energy (ACE) and gliding values (Rakesh et al., 2015, 2016; Satapathy et al., 2020).

#### Statistical analysis

All data were expressed as mean ± standard deviation (*n* = 3). Results were determined using one-way analysis of variance (ANOVA), followed by Duncan’s multiple range test using GraphPad Software, Inc (version 6.0, California, USA). The results were considered statistically significant if the *P* < 0.05. The minimum dosage of extract that is necessary to produce 50% inhibition was known as the effective dose (ED_50_), which is calculated using regression analysis.

## Results & Discussion

Phytofractions are extensively used as nutraceuticals in complementary and alternative medicine to boost health and to prevent stress, inflammation, and secondary lifestyle diseases. The current study investigates the interconnection of antioxidants and regulation of calcium oxalate which is a burgeoning area of research that is gaining increased attention due to the increased number of urolithiasis cases globally. We examined the ability of naturally occurring plant-derived aqueous fractions for their anti-stress and anti-urolithiatic efficacy in the prevention and treatment of the disease using *in vitro* and *in silico* platforms.

With the background of Indian traditional medicine (Ayurveda) and folklore, twenty priority botanicals with various other bioactivities were selected namely *Aegle marmelos, Apium graveolens, Artocarpus heterophyllus, Azadirachta indica, Citrus limon, Citrus sinensis, Coriandrum sativum, Cucumis sativus, Kalanchoe pinnata, Lawsonia inermis, Mangifera indica, Manilkara zapota, Momordica charantia, Moringa oleifera, Musa paradisiaca, Phyllanthus emblica, Plectranthus amboinicus, Psidium guajava, Punica granatum* and *Ribes Uva-crispa* (table 1) and their leaves component were screened for their antioxidant and anti-urolithiatic potency using aqueous hot extraction method. This extraction method was intentionally used with the purpose to formulate homemade decoctions with the motto of making food as medicine. The phyto-cocktails from all 20 extracts were subjected to qualitative phytochemical profiling for the presence of carbohydrates, glycosides, alkaloids, phenols, flavonoids, tannins, triterpenoids, steroids, saponin, and gums (supplementary table 1). Among them, most of the aqueous extract was found to be composed of the abovementioned phytochemical classes. However, glycosides and steroids were absent in most of the extracts indicating the non-dissolution of hydrophobic phyto-molecules. Most of the hydrophilic phyto-molecules (tight and loosely bound secondary metabolites) were extracted during the procedure. This observation implies an increased bioavailability of the phytococktail with enhanced bioabsorption within the system. All aqueous extracts were further subjected to the estimation of total phenolic content (TPC) to ensure the quantitative measurements (figure 1). Among the 20 leaves extracts *P.granatum, M.oleifera, P.guajava,* and *C.sativum* contented higher levels (nearing 2 mg/mL) whereas, *M.charantia* and *C.sativus* exhibited the least (nearing 0.5 mg/mL) phenolic content. The rest of the 14 extracts possessed (a range between1 to 1.5 mg/mL) of phenolics. For all dose-dependent studies, TPC was considered for quantitation.

**Figure 1:**
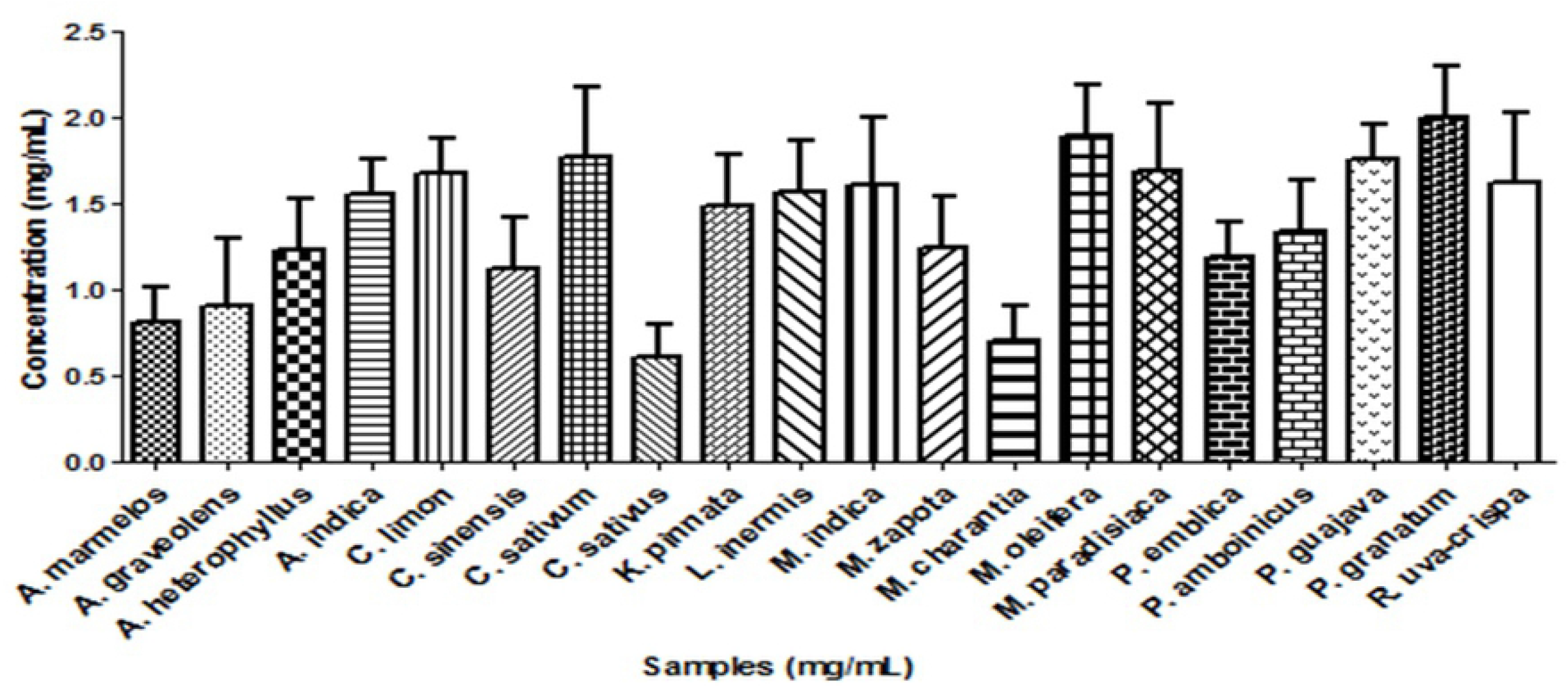
Quantitative estimation of polyphenolic content of 20 selected botanicals.

Oxyradicals majorly reactive oxygen species (ROS) and reactive nitrogen species (RNS) are generated during metabolic insult and are the prime cause for biomolecular dysregulation, cellular damage, accounting for cell membrane disruption, mitochondrial dysbiosis, protein folding, inflammation leading to a high frequency of time-dependent diseases like early aging, cancer and neurodegenerative disorders which hampers general health maintenance. According to Vina and co-workers (2005), oxidants are the primordial cause for any disorders. Striking a proper equilibrium with antioxidant molecules will largely facilitate the chances of better survival. Hence a quest for exploring potent phyto-antioxidants is an urgent necessity to overcome lifestyle-related diseases (Garg et al., 2020). In the current study, all the 20 aqueous hot leaf extracts were subjected for their efficacy to quench free radical scavenging (figure 2) in a dose-dependent manner at 5μg/mL, 15μg/mL, and 25μg/mL of TPC respectively. DPPH (figure 2A), metal chelating (figure 2B), and reducing power (figure 2C) assays were employed. Ethylenediaminetetraacetic acid (EDTA) was used as a standard for metal chelating assay and gallic acid was used as the standard for the other two assays. In DPPH assay, *A.indica, K.pinnata, P.granatum,* and *P.emblica* were found to have better free radical scavenging activity. Whereas, *A.graveolens, C.sativum, C.sativus,* and *P.amboinicus* indicated the least activity among the 20 screened botanicals (figure 2A). However, *M.zapota, M.charantia,* and *M.oleifera* exhibited better metal chelating ability. Further, *A.indica* and *C.sativus* showed the least activity among the screened phytoextracts (figure 2B). The reducing power assay was well responded with *C.limon, K.pinnata, L. inermis, M.indica, P.emblica,* and *P.granatum* extracts and on the contrary, *A.heterophyllus, C.sativus, M.charantia, P.amboinicus,* and *P.guajava* exhibited the least activity (figure 2C). Similar results (Bidchol et al., 2011; Wong et al., 2014; Nedamani et al., 2015) have been well document by the previous researchers.

**Figure 2:**
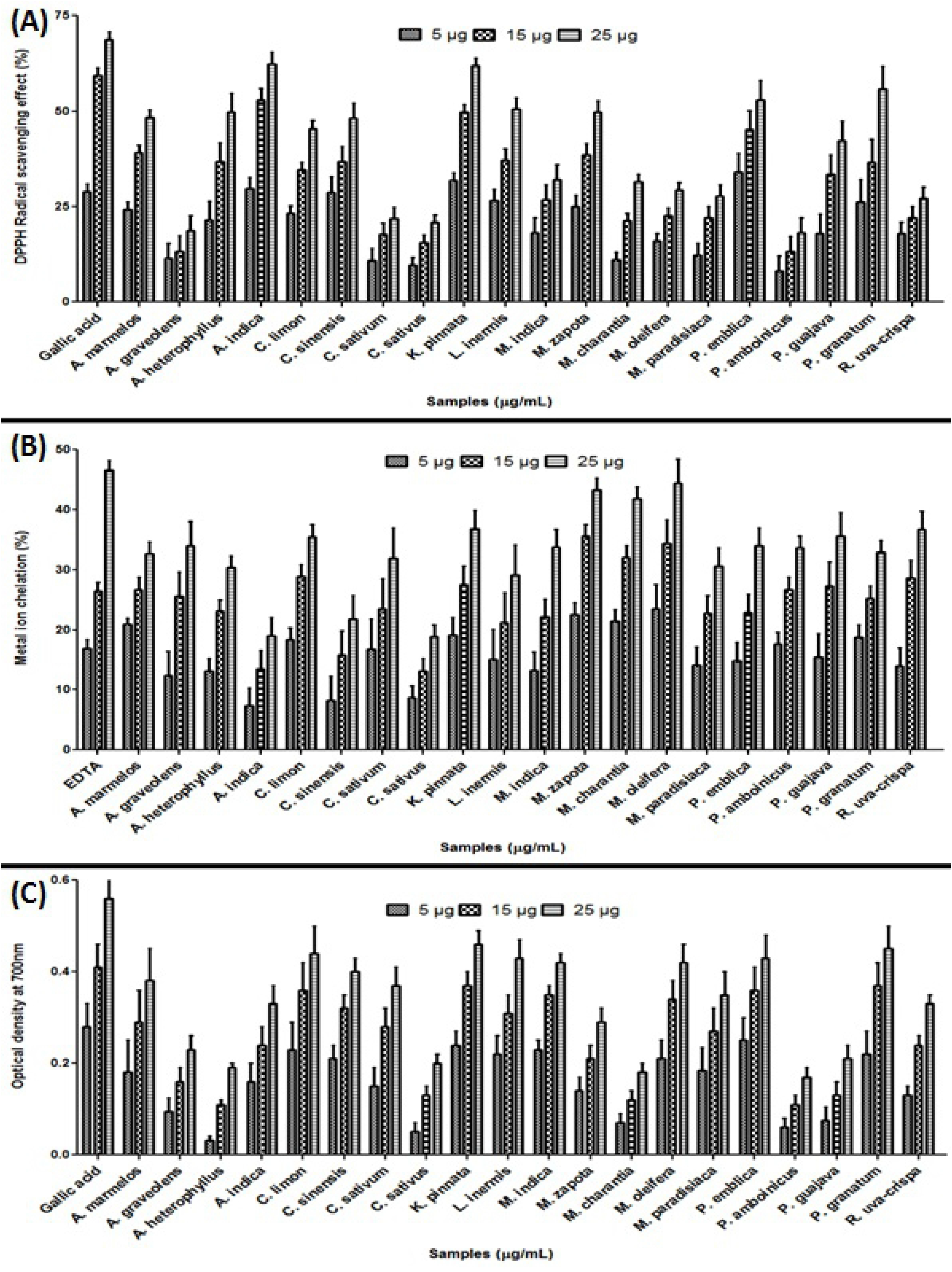
Antioxidant radical scavenging assay of the 20 selected botanicals at 5μg/mL, 15 μg/mL and 25 μg/mL respectively. (A): DPPH assay, (B): Metal ion chelating assay and (C): Reducing power assay.

Besides, the antioxidant potency, the 20 selected botanicals with a concentration of 25μg/mL were subjected to spectral studies, to assess their ability in inhibiting calcium oxalate crystal formation with cystone as the standard herbal drug (figure 3). The results exhibited that among the screened extracts *C.sativum, K.pinnata, M.oleifera,* and *M. paradisiaca,* were found to have evidence in inhibiting calcium oxalate crystal formation. Henceforth for all further studies, these five aqueous phytoextracts were considered. Further, the top five extracts were subjected to an *in vitro* experimental chicken egg membrane model to mimic the dissolution across the biological membrane (figure 4) of which *K.pinnata* exhibited 71% dissolution than other extracts in comparison to cystone standard (86%). Hence for all the further experimentation leaves aqueous hot extract of *K.pinnata* was used and subjected to critical analyses.

**Figure 3:**
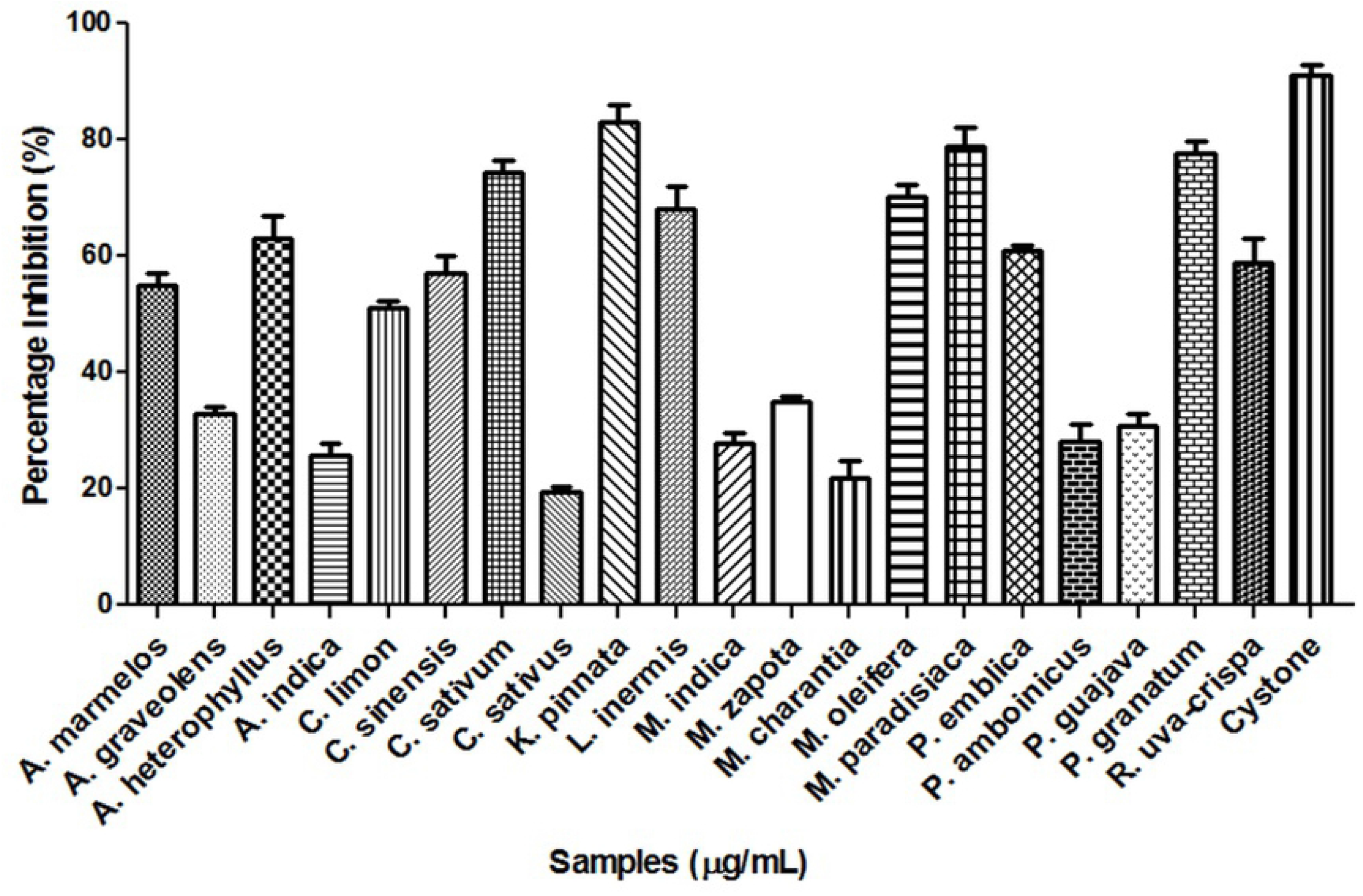
Anti-lithiatic activity of 20 botanicals

**Figure 4:**
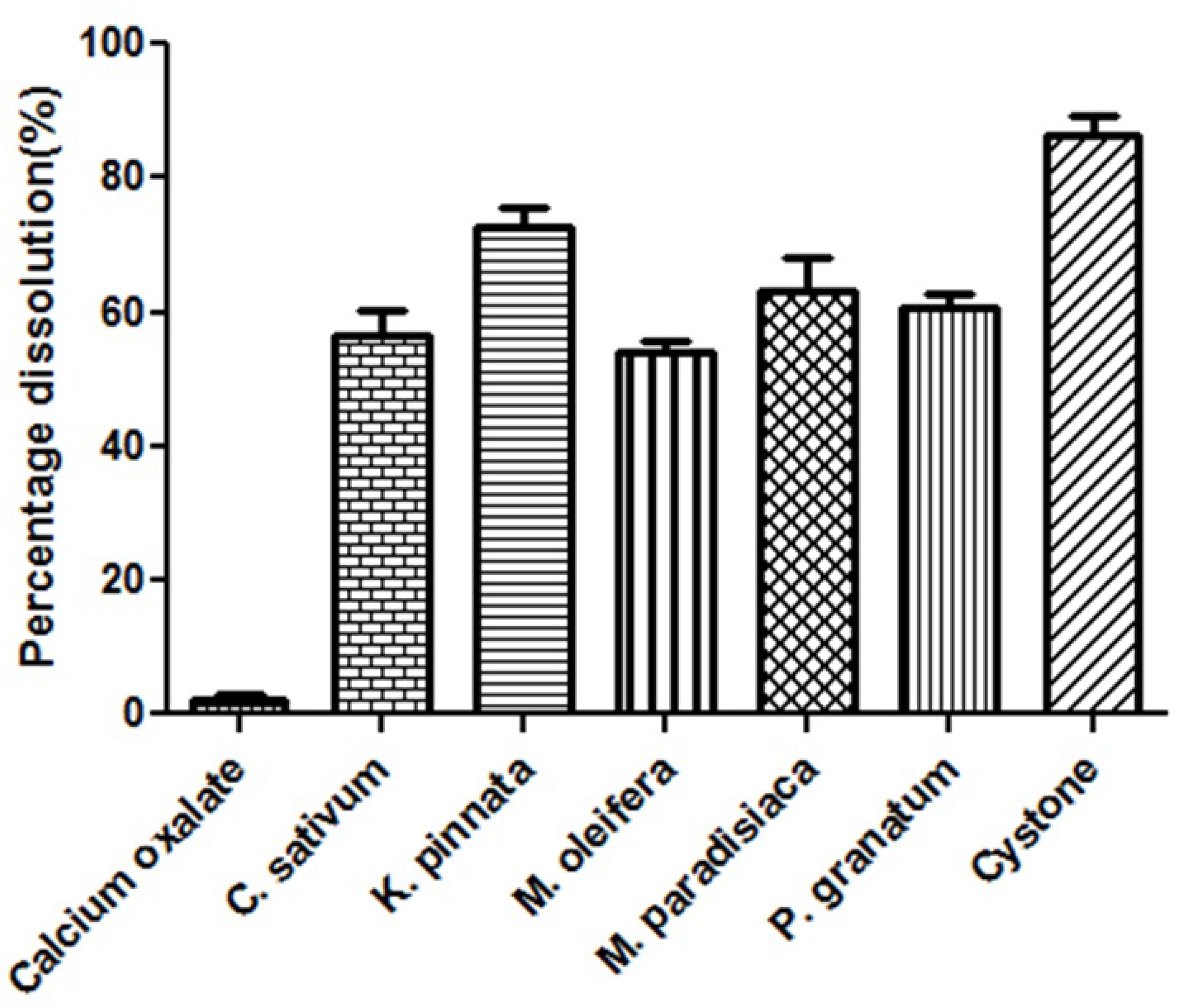
In vitro experimental assessment of anti-lithiatic activity of top 5 best phyto-contributors using chicken egg membrane model with cystone as standard herbal drug.

Nucleation and aggregation are the two major phases during lithiasis. To assess the sequential effectiveness of *K.pinnata*, the extract was subjected in a dose-dependent manner at a range of concentrations (5μg/mL, 10μg/mL, 15μg/mL, 20μg/mL, and 25μg/mL) respectively with 25μg/mL of cystone as standard drug formulation. The calcium oxalate curve was considered as untreated standard (supplementary figure 1) The nucleation assay was assessed at 1h, 3h, and 24 h after incubation (figure 5A). The results provided sufficient evidence that after 24 h of incubation a promising inhibition (70%) of calcium oxalate was observed with *K.pinnata* extract compared to cystone (76%). Further, aggregation assessment was also performed to have a microscopic birds’ eye view on the crystal dissolution (figure 5B). Thus, *K.pinnata* extract was found to be a potent anti-crystallization contributor. This could be probably due to higher concentrations of flavonoids, saponins, and gums (Van-Dooren et al., 2016) which has a shred of qualitative evidence with phytochemical profiling (supplementary table 1).

**Figure 5:**
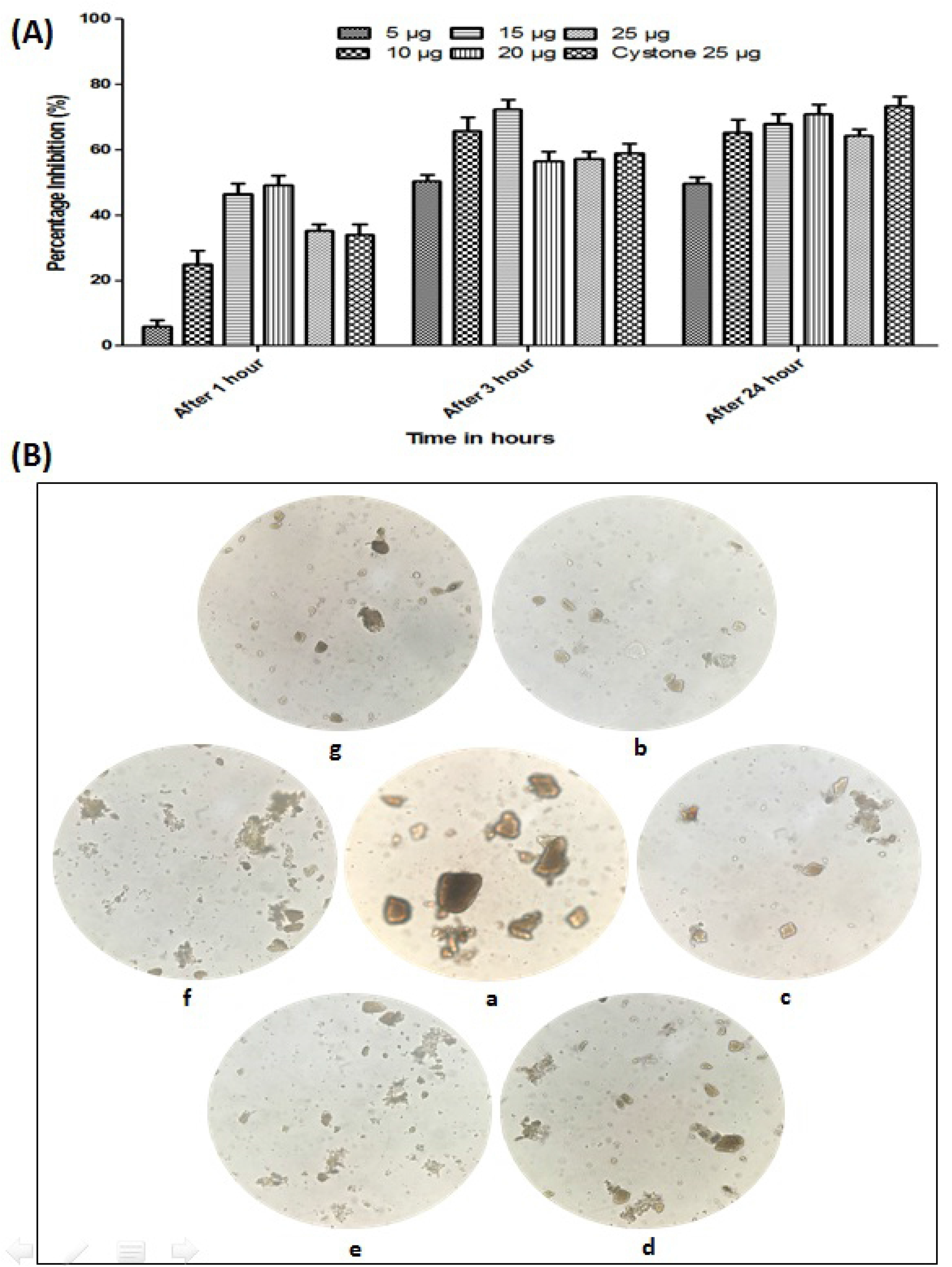
**(A):** Dose-dependent assessment of KP extract using nucleation assay. (B): Dose-dependent assessment of KP extract using aggregation assay.

Further, to evaluate the toxicity of the *K.pinnata* extract RBC (figure 6: Panel A) and DNA (figure 6: Panel B) protection assay were performed. The *K.pinnata* phyto-cocktail exhibited potential hampering of the induced oxidant partially at 5μg/mL and completely at 25μg/mL in comparison to gallic acid (pure drug) and cystone (formulation) as standard drug. Similar kind of studies has been evident with Beulah et al., (2015) and Meghashri et al., (2010) respectively. HPLC profiling was performed (figure 7) on aqueous *K.pinnata* leaves extract. The sample was found to contain gallic acid at 272nm, rutin at 257nm, and vanillic acid at 261nm whereas, ferulic acid and quercetin were present in minor quantities. The chromatogram for standards has been provided in supplementary figure 2.

**Figure 6:**
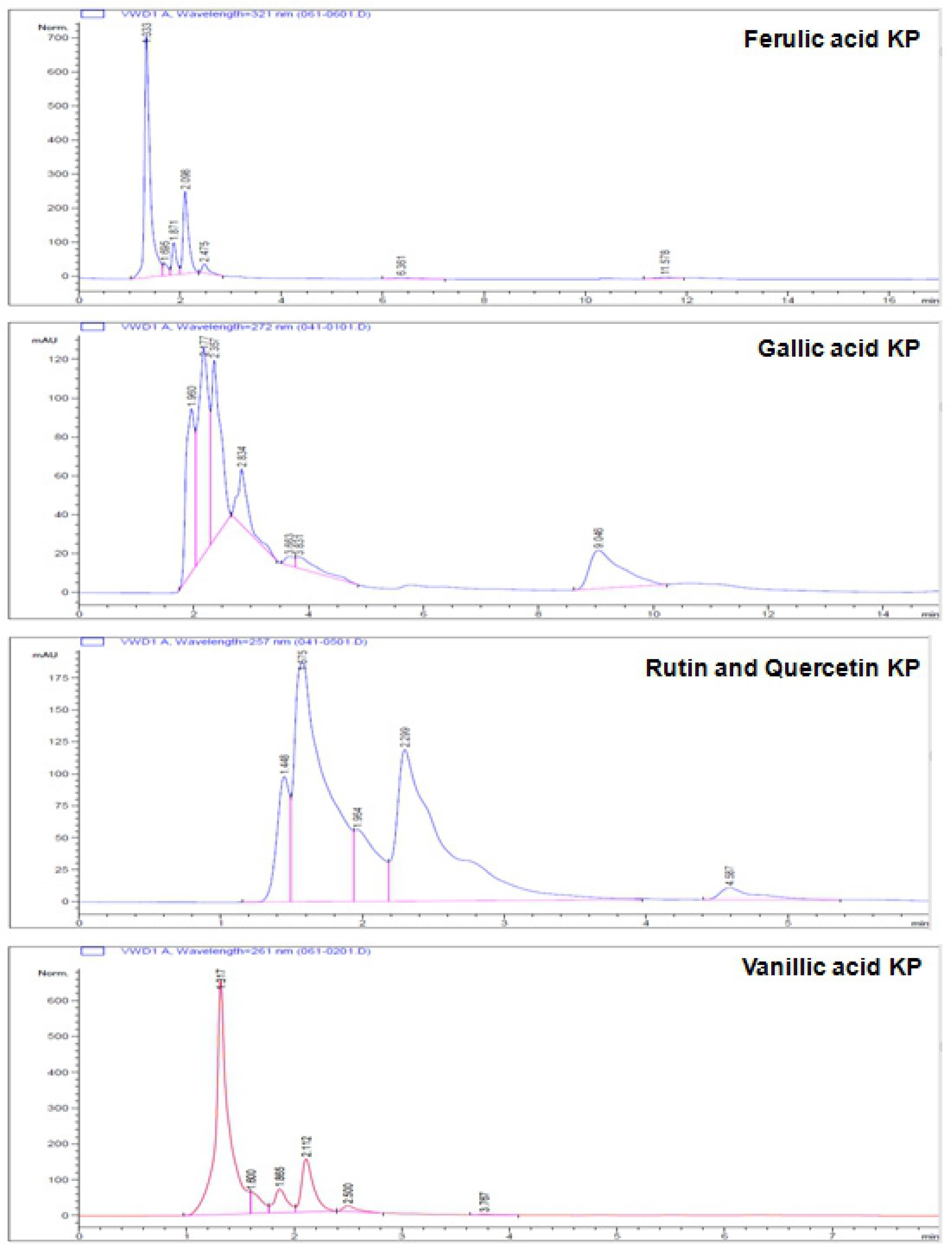
Panel A: Evaluation of KP extracts on erythrocyte morphology by RBC protection assay, (a): control-RBC, (b): RBC+PBS (pH 7.4), (c): RBC plus oxidant, (d): RBC plus gallic acid (std 10μg/mL), (e): RBC plus 25μg/mL of KP and oxidant after lh of incubation. Panel B: Electrophoretic analysis of DNA protection assay by KP extracts. Lane 1: native DNA, Lane 2: oxidised DNA, Lane 3: Gallic acid (std 10μg/mL), Lane 4: Cystone (std 10μg/mL), Lane 5: KP extract (5μg/mL) and Lane 6: KP extract (25μg/mL).

**Figure 7:**
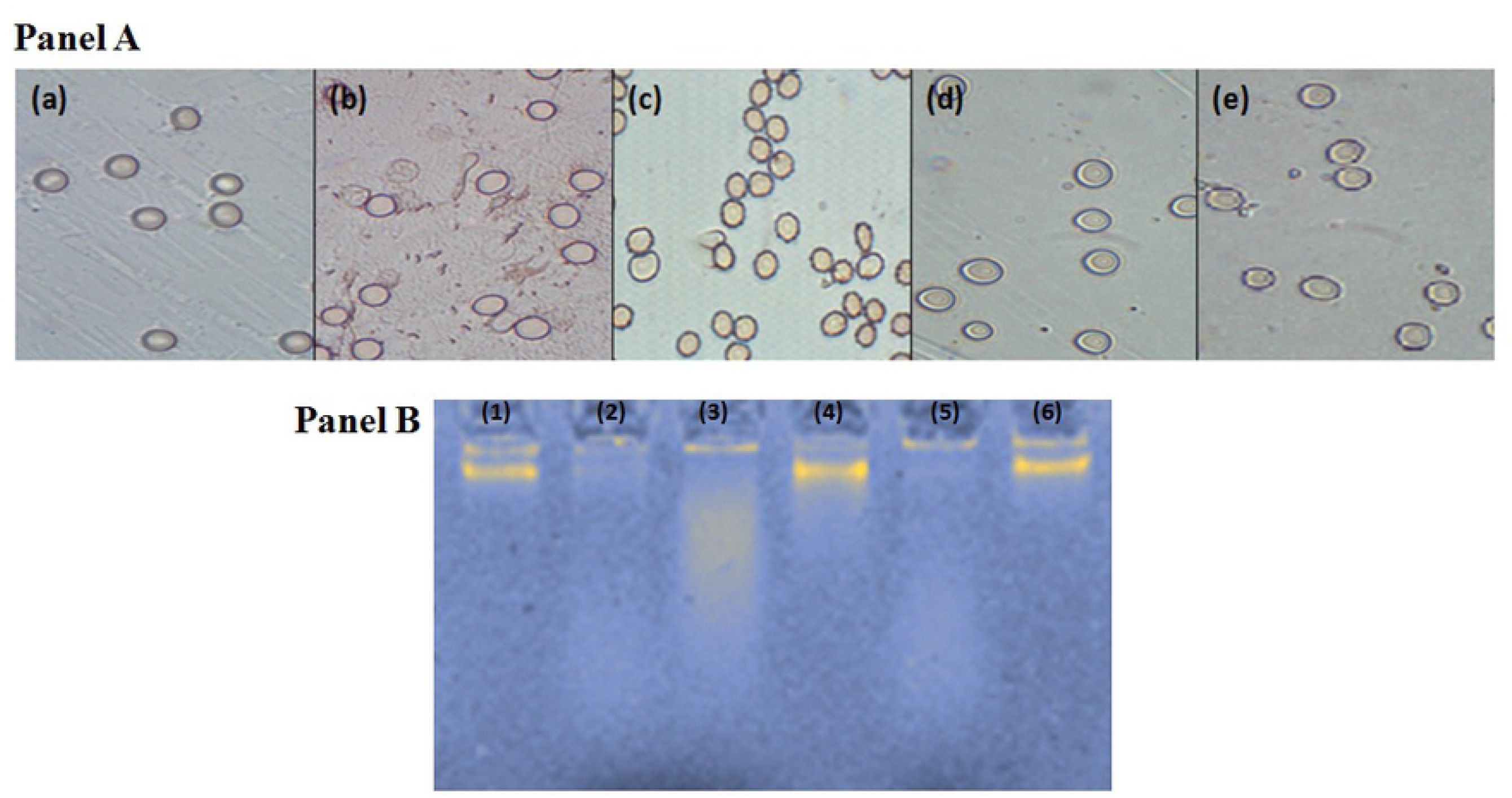
HPLC profiling of the aqueous KP extract containing the phytochemicals.

Inside a cell, biomarkers such as antioxidant, inflammatory, and calcium oxalate regulating metabolic enzymes play a pivitol role in maintaining cellular homeostasis during lithiasis. Hence an attempt was made to investigate these two classes of enzymes namely catalase (CAT), superoxide dismutase (SOD), peroxidase (PER), glutathione S-transferase (GST), contributing towards redox machinery and stress respectively. Besides, metabolic enzymes such as alanine-glyoxlate aminotransferase, oxalyl-coA decarboxylase, D-glycerate dehydrogenase, and lactate dehydrogenase (LDH) regulate a crucial role in stone formation, and inflammation was considered for the study. Three major ligands namely, gallic acid, rutin, and vanillic acid were considered with potassium-magnesium citrate as the standard-currently used drug for urolithiasis. Structure-activity relationship (SAR) studies were performed for antioxidant (figure 8) and calcium oxalate regulating enzymes (figure 9) with the best-docked pose. Further, the data have been tabulated in table 2 and table 3 respectively indicating the specific amino acid which comprises the active site for the target and ligand. Among the docked ligands rutin (molar mass: 610.517 g/mol) was found to be active and best docked on both the classes of the target enzymes than compared to the available drug currently used by clinicians to treat uro/nephrolithiasis. Previous studies on rutin as a potent antioxidant and anti-urolithiatic agent have been well document by Enogieru, et al., (2018) and Ghodasara et al., (2010) respectively. Further, Radwan et al (2021) and Santos et al., (2021) have explored the possible role of the *K. pinnata* attributes in hemostasis and oxytocin signaling pathways respectively.

**Figure 8:**
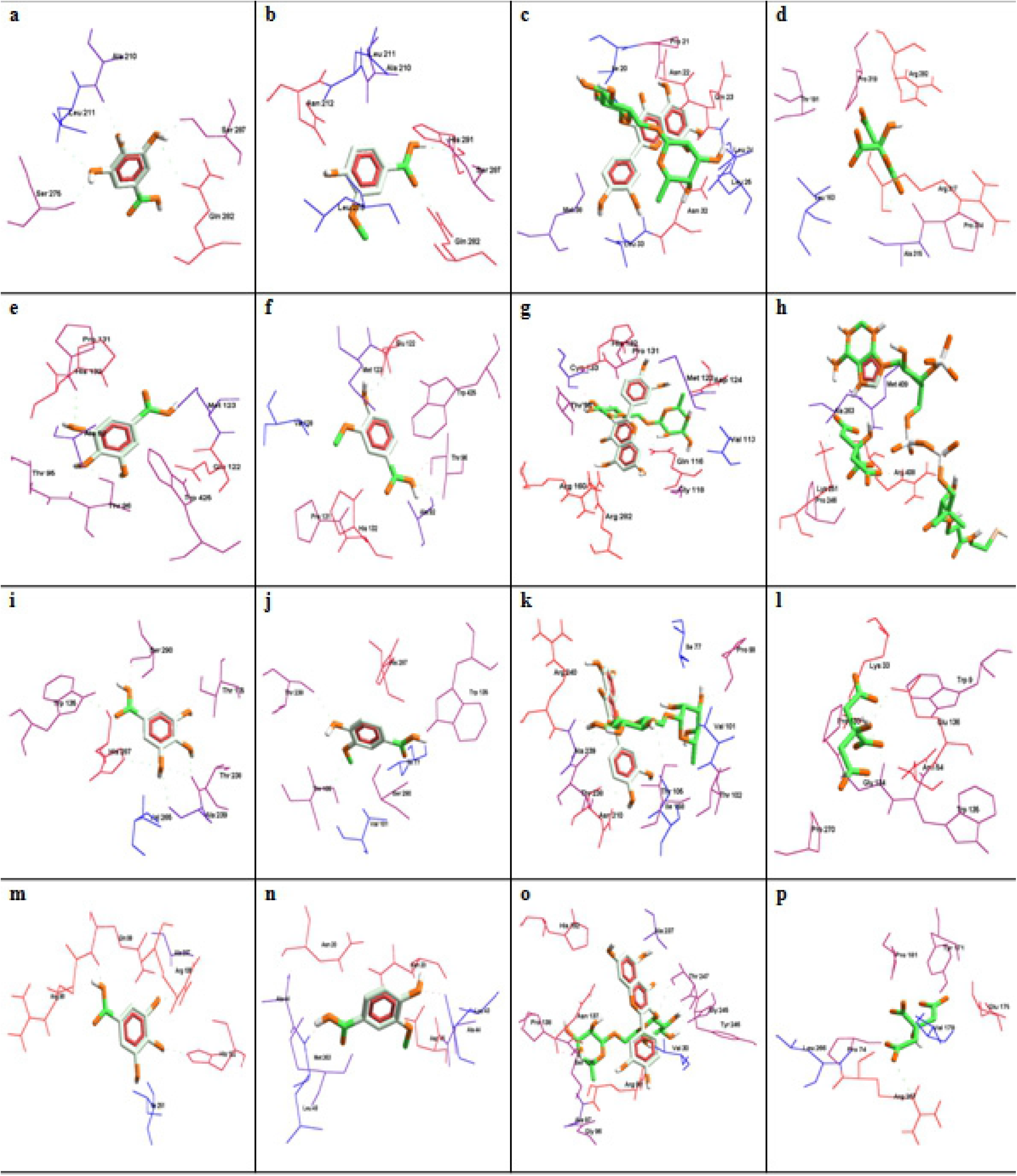
*In silico* molecular docking of KP phytochemical ligands on antioxidant enzymes. **Footnotes for Figure 8:** Docked images of antioxidant enzymes: (a) Superoxide dismutase docked with Gallic acid, (b) Superoxide dismutase docked with Vanillic acid, (c) Superoxide dismutase docked with Rutin, (d) Superoxide dismutase docked with Potassium-Magnesium Citrate, (c) Peroxidase docked with Gallic acid, (f) Peroxidase docked with Vanillic acid, (g) Peroxidase docked with Rutin, (h) Peroxidase docked with Potassium-Magnesium Citrate, (i) Catalase docked with Gallic acid, (j) Catalase docked with Vanillic acid, (k) Catalase docked with Rutin, (I) Catalase docked with Potassium-Magnesium Citrate, (m) Glutathione S-transferase docked with Gallic acid, (n) Glutathione S-transferase docked with Vanillic acid, (o) Glutathione S-transferase docked with Rutin and, (p) Glutathione S-transferase docked with Potassium-Magnesium Citrate.

**Figure 9:**
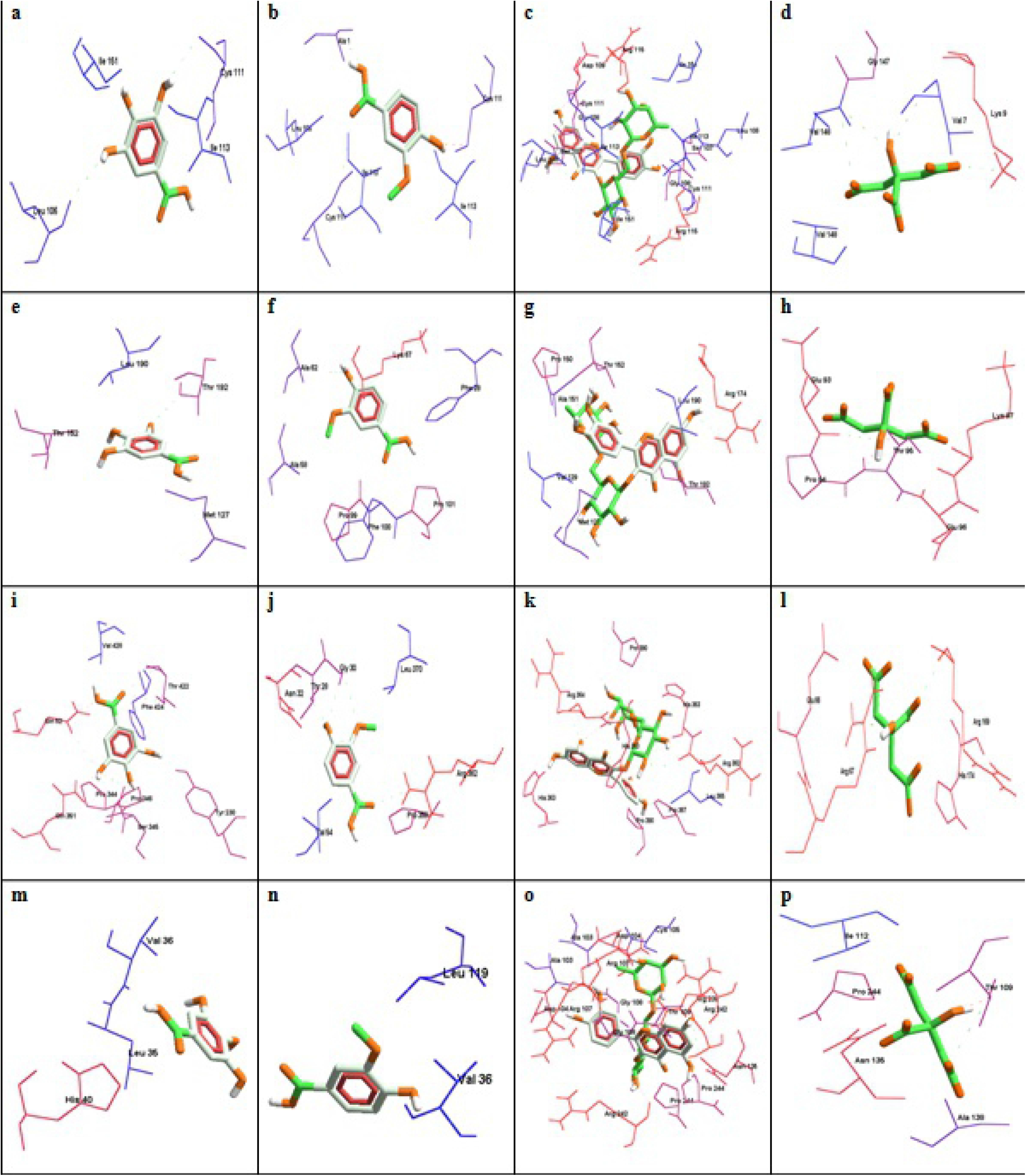
*In silico* molecular docking of KP phytochemical ligands on calcium oxalate regulating metabolic enzymes. **Footnotes for Figure 9:** Docked images of metabolic enzymes: (a) Alanine glyoxylate aminotransferase docked with Gallic acid, (b) Alanine glyoxylate aminotransferase docked with Vanillic acid, (c) Alanine glyoxylate aminotransferase docked with Rutin, (d) Alanine glyoxylate aminotransferase docked with Potassium-Magnesium Citrate, (c) Oxalyl CoA decarboxylase docked with Gallic acid, (f) Oxalyl CoA decarboxylase docked with Vanillic acid, (g) Oxalyl CoA decarboxylase docked with Rutin, (h) Oxalyl CoA decarboxylase docked with Potassium-Magnesium Citrate, (i) D- glycerate dehydrogenase docked with Gallic acid, (j) D- glycerate dehydrogenase docked with Vanillic acid, (k) D- glycerate dehydrogenase docked with Rutin, (1) D- glycerate dehydrogenase docked with Potassium-Magnesium Citrate, (m) Lactate dehydrogenase docked with Gallic acid, (n) Lactate dehydrogenase docked with Vanillic acid, (o) Lactate dehydrogenase docked with Rutin and, (p) Lactate dehydrogenase docked with Potassium-Magnesium Citrate.

**Table 2:** Structure-activity relationship (SAR) of potent ligands from KP with antioxidant enzymes.

| Name of the Antioxidant enzymes (Targets) | Name of the molecules (Ligands) | Details of H-bond interaction |  |  | Atomic Contact Energy (ACE) values in KJ/mol | Gliding values | Amino acid residues of docked domains |
| --- | --- | --- | --- | --- | --- | --- | --- |
|  |  | No. of bond | Bond energy | Bond length |  |  |  |
| Superoxide dismutase | Gallic acid | 2 | -2.35 | 3.12 | -137.22 | -2.74 | Leu 106, Ile 151, Cys 111, Ile 113 |
|  | Vanillic acid | 4 | -2.5 | 2.60 | -156.44 | -3.12 | Ala 1, Leu 106, Cys 111, Ile 113, Cys 111, Ile 113 |
|  | Rutin | 10 | -2.36 | 2.60 | -349.88 | -6.99 | Leu 106, Ser 107, Gly 108, Cys 111, Ile 113, Arg 115, Ile 151, Leu 106, Ser 107, Gly 108, Asp 109, Cys 111, Ile 113, Arg 115, Ile 151 |
|  | Potassium Magnesium Citrate (STD) | 4 | -2.5 | 2.65 | -222.84 | -4.45 | Val 148, Val 7, Lys 9, Gly 147, Val 148 |
| Peroxidase | Gallic acid | 1 | -2.5 | 2.90 | -89.50 | -1.79 | Met 127, Thr 152, Leu 190, Thr 192 |
|  | Vanillic acid | 1 | - | - | -116.04 | -2.32 | Phe 28, Ala 58, Ala 62, Lys 67, Pro 99, Phe 100, Pro 101 |
|  | Rutin | 8 | -2.04<br>-2.5<br>-2.5 | 3.19<br>2.63<br>2.68 | -243.50 | -4.87 | Arg 174, Leu 190, Thr 192, Met 127, Val 129, Pro 150, Ala 151, Thr 152 |
|  | Potassium Magnesium Citrate (STD) | 4 | -2.5 | 3.05 | -184.76 | -3.69 | Glu 93, Pro 94, Thr 95, Glu 96, Lys 97 |
| Catalase | Gallic acid | 9 | -2.05<br>-2.26<br>-2.5 | 3.18<br>3.14<br>2.88 | -53.17 | -1.06 | Gln 52, Tyr 230, Pro 344, Ser 345, Pro 346, Gln 351, Thr 422, Phe 424, Val 428 |
|  | Vanillic acid | 5 | -2.06<br>-2.5 | 2.54<br>2.87 | -53.80 | -1.07 | Pro 358, Arg 362, Leu 370, Thr 28, Gly 30, Asn 32, Val 54 |
|  | Rutin | 8 | -2.5<br>-2.5<br>-2.27<br>-2.43 | 2.82<br>2.63<br>3.14<br>2.79 | -217.45 | -4.34 | His 363, Pro 390, Arg 362, His 363, Leu 365, Pro 367, His 363, Arg 364, Pro 390 |
|  | Potassium Magnesium Citrate (STD) | 2 | -1.98 | 3.20 | -78.17 | -1.56 | Glu 66, Arg 67, Arg 169, His 174 |
| Glutathione S-transferase | Gallic acid | - | - | - | -165.42 | -3.30 | Leu 35, Val 36, His 40 |
|  | Vanillic acid | - | - | - | -176.76 | -3.53 | Val 36, Leu 119 |
|  | Rutin | 12 | -2.40<br>-2.5<br>-2.5 | 3.11<br>2.91<br>3.07 | -196.14 | -3.92 | Ala 103, Asp 104, Arg 107, Gly 108, Arg 239, Arg 242, Pro 244, Ala 103, Asp 104, Cys 105, Arg 107, Gly 108, Thr 109, Asn 135, Arg 242, Pro 244 |
|  | Potassium Magnesium Citrate (STD) | 2 | - | - | -251.96 | -5.03 | Thr 109, Ile 112, Asn 135, Ala 138, Pro 244 |

**Table 3:** Structure-activity relationship (SAR) of potent ligands from KP with calcium regulating metabolic enzymes.

| Name of the Metabolic enzymes (Targets) | Name of the molecules (Ligands) | Details of H-bond interaction |  |  | Atomic Contact Energy (ACE) values KJ/mol | Gliding Values | Amino acid residues of docked domains |
| --- | --- | --- | --- | --- | --- | --- | --- |
|  |  | No. of bond | Bond energy | Bond length |  |  |  |
| Alanine-Glyoxylate Aminotransferase | Gallic acid | 5 | -2.5<br>-2.5<br>-2.5<br>-2.5 | 2.87<br>3.08<br>2.68<br>2.76 | -110.58 | -2.21 | Ala 210, Leu 211, Ser 275, Gln 282, Ser 287 |
|  | Vanillic acid | 2 | -2.5<br>-2.5 | 2.99<br>2.68 | -134.52 | -2.69 | Ala 210, Leu 211, Asn 212, Leu 278, Gln 282, Ser 287, His 291 |
|  | Rutin | 3 | -2.25<br>-2.10 | 3.14<br>3.17 | -302.13 | -6.04 | Ile 20, Pro 21, Asn 22, Gln 23, Leu 24, Leu 25, Asn 32, Leu 33, Met 38 |
|  | Potassium Magnesium Citrate (STD) | 1 | - | - | -202.38 | -4.04 | Leu 163, Thr 191, Arg 292, Pro 314, Ala 315, Arg 317, Pro 319 |
| Oxalyl-CoA decarboxylase | Gallic acid | 5 | -2.5<br>-2.5<br>-2.09 | 2.86<br>2.70<br>2.78 | -121.82 | -2.43 | Ala 92, Thr 95, Thr 96, Pro 131, His 132, Trp 425, Glu 122, Met 123 |
|  | Vanillic acid | 4 | -2.34 | 2.81 | -130.17 | -2.60 | Ala 92, Thr 96, Pro 131, His 132, Trp 425, Glu 122, Met 123, Val 128 |
|  | Rutin | 8 | -2.39<br>-2.06 | 3.12<br>3.18 | -211.63 | -4.23 | Thr 95, Pro 131, His 132, Cys 133, Arg 160, Arg 282, Val 113, Gln 116, Gly 118, Met 123, Asp 124 |
|  | Potassium Magnesium Citrate (STD) | 3 | -2.05 | 3.18 | -154.10 | -3.08 | Pro 246, Lys 251, Ala 263, Arg 408, Met 409 |
| D-glycerate dehydrogenase | Gallic acid | 7 | -2.35<br>-2.5<br>-2.5<br>-2.09 | 2.58<br>3.08<br>3.04<br>2.55 | -83.16 | -1.66 | Thr 105, Thr 238, Ala 239, Val 265, His 287, Ser 290, Trp 135 |
|  | Vanillic acid | 3 | -2.20 | 2.56 | -90.91 | -1.81 | Trp 135, Ile 77, Val 101, Thr 105, Thr 238, His 287, Ser 290 |
|  | Rutin | 7 | -2.45<br>-2.17 | 3.10<br>3.16 | -261 | -5.22 | Ile 77, Pro 98, Val 101, Thr 102, Thr 105, Ile 158, Asn 210, Thr 238, Ala 239, Arg 240 |
|  | Potassium Magnesium Citrate (STD) | 5 | -2.5<br>-2.5 | 2.90<br>2.78 | -105.07 | -2.10 | Pro 133, Gly 134, Trp 135, Glu 136, Trp 9, Lys 33, Asn 54, Pro 270 |
| Lactate dehydrogenase | Gallic acid | 3 | -2.02 | 3.19 | -43.36 | -0.86 | Arg 98, Gln 99, Arg 105, His 192, Ala 237, Ile 251 |
|  | Vanillic acid | 2 | - | - | -61.57 | -1.23 | Asn 20, Leu 43, Ala 44, Asp 45, Asn 20, Leu 43, Ala 44, Met 263 |
|  | Rutin | 11 | -2.5<br>-2.5 | 2.71<br>2.66 | -250.63 | -5.01 | Val 30, Gly 96, Ala 97, Arg 98, Ser 136, Asn 137, Pro138, His 192, Ala 237, Gly 245, Tyr 246, Thr 247 |
|  | Potassium Magnesium Citrate (STD) | 3 | -2.5 | 2.91 | -109.89 | -2.19 | Leu 266, Arg 267, Pro 74, Tyr 171, Glu 175, Val 179, Pro 181 |

Finally, to summarize, the current study has made an exciting observation that *K.pinnata* aqueous leaves extract can completely suppress the clinical disease in an experimental *in vitro* model of urolithiasis. However, further studies at a molecular level are needed to test the central hypothesis that *K. pinnata* has a profound effect on pro- and antiinflammatory responses regulating transcriptional activity which may lead to the amelioration of the disease. Newer in silico algorithms have to be generated which can target gene prediction and pathways that are dysregulated during urolithiasis. In conclusion, these studies should provide novel insights into the basic mechanisms through which phyto-cocktails exhibits anti-lithiatic properties so that it can be used against a wide range of lifestyle diseases leading to better health and wellbeing.

## Acknowledgments

Ms. Aishwarya Tripurasundari Devi would like to thank the Indian Council of Medical Research (ICMR) for the award of Senior Research Fellow (SRF) Award Letter Number - File no.5/3/8/55/ITR-F/2018-ITR dated 11.6.2018. The authors thank, Dr. Trevani, Herbarium Section, Department of Studies in Botany, University of Mysore, for identification of the screened botanicals. All authors thank the Hon’ble Vice-Chancellor, JSS Science and Technology University, Principal, SJCE, Mysore for his encouragement, and JSS research foundation for their constant inspiration. All the authors are thankful to Dr. Shaukath Ara Khanum, Department of Chemistry, Yuvaraja College, and Dr. Shubha Gopal, Department of Studies in Microbiology, University of Mysore, Mysore for the long-term collaboration and guidance in understanding the biology of phytochemical molecules. Further, we extend our gratitude towards the management and office bearers of Dayananda Sagar University (DSU), Bengaluru, Karnataka, India, for constant inspiration, motivation, and encouragement to pursue scientific research. Special thanks to Prof. Sunil S. More, Dean, School of Basic and Applied Sciences, DSU for all approval and permissions during the execution of this research work.

